# TRTools: a toolkit for genome-wide analysis of tandem repeats

**DOI:** 10.1101/2020.03.17.996033

**Authors:** Nima Mousavi, Jonathan Margoliash, Neha Pusarla, Shubham Saini, Richard Yanicky, Melissa Gymrek

**Affiliations:** Department of Electrical and Computer Engineering, University of California San Diego, 9500 Gilman Drive, MC 0639, La Jolla, CA, 92093, USA; Department of Medicine, University of California San Diego, 9500 Gilman Drive, MC 0639, La Jolla, CA, 92093, USA; Department of Bioengineering, University of California San Diego, 9500 Gilman Drive, MC 0639, La Jolla, CA, 92093, USA; Department of Computer Science and Engineering, University of California San Diego, 9500 Gilman Drive, MC 0639, La Jolla, CA, 92093, USA

## Abstract

**Summary:** A rich set of tools have recently been developed for performing genome-wide genotyping of tandem repeats (TRs). However, standardized tools for downstream analysis of these results are lacking. To facilitate TR analysis applications, we present TRTools, a Python library and a suite of command-line tools for filtering, merging, and quality control of TR genotype files. TRTools utilizes an internal harmonization module making it compatible with outputs from a wide range of TR genotypers.

**Availability:** TRTools is freely available at https://github.com/gymreklab/TRTools.

**Supplementary information:** Supplementary data are available at *bioRxiv*.

## 1 Introduction

Tandem repeats (TRs) represent one of the largest sources of human genetic variation and are well-known to affect many human phenotypes [4]. Improvements in sequencing technology and bioinformatics algorithms have led to the recent development of a rich set of tools for performing genome-wide analysis of TR variation [8, 7, 5, 3, 1]. These tools take aligned sequencing reads as input and output Variant Call Format (VCF) files containing estimates of TR copy number at one or more genomic TRs. The resulting VCF files may be used for a wide variety of downstream applications. However, before doing so it is usually necessary to perform filtering, quality control (QC), and merging of files across samples. While utilities exist for performing such manipulations on VCF files containing SNP variants, these tools often do not handle multi-allelic TRs and are not designed to compute TR-specific statistics. Further, different TR genotypers use different allele annotations, complicating the use of downstream tools.

Here, we present TRTools, an open-source toolkit for performing analyses on TR genotypes. TRTools provides utilities for filtering, merging, comparing, and performing QC on TR VCF files. It is compatible with five major TR genotypers (GangSTR, HipSTR, ExpansionHunter, PopSTR, and adVNTR) and can easily be extended to handle VCFs from additional tools.

## 2 Features and Methods

TRTools consists of a suite of command-line utilities and a corresponding Python library for performing common operations on TR genotypes, including filtering, callset comparisons, and other workflows. It parses VCF files using the PyVCF [2] library and implements a “TR harmonizer” module that converts VCF formats from each tool to a standardized representation. This harmonization step enables downstream operations to proceed agnostic of the original tool used to produce the genotypes. For all utilities described below, the `--vcftype` argument may be used to specify the genotyping tool used. If not specified, the type is automatically inferred. In the following sections, we summarize the current functionality available in TRTools. Utilities are summarized in Table 2.2.

### 2.1 DumpSTR

`dumpSTR` is a tool for filtering TR VCF files. It performs call-level filtering (e.g., minimum call depth, maximum TR stutter error) and locus-level filtering (e.g., minimum call rate or deviation from Hardy-Weinberg Equilibrium). `dumpSTR` is specially built to handle VCF FORMAT and INFO fields unique to TR genotypers. Unlike standard VCF filtering tools, it also computes locus-level metrics such as heterozygosity and Hardy-Weinberg Equilibrium based on TR allele lengths. It takes as input a VCF file and outputs a new VCF with locus-level filters annotated in the FILTER column and call-level filters annotated in the FORMAT field for each call.

~~~
dumpSTR --vcf VCF --out OUTPREFIX \
   [--vcftype={eh|gangstr|hipstr|popstr|advntr}] \
   [filters]
~~~

### 2.2 MergeSTR

`mergeSTR` is a method for merging VCF tools generated by TR genotyping methods. While methods for merging VCF files currently exist [6], TR VCFs have unique characteristics that call for a specialized merging tool. TRs are often multi-allelic, and VCFs called on different sample sets may contain different alternate allele sets. Further, TR alleles may be trimmed to remove redundant TR sequence, which can interfere with downstream analysis of TR lengths. `mergeSTR` takes two or more VCFs as input and outputs a merged VCF file containing all of the samples included in the input VCFs.

~~~
mergeSTR --vcfs VCF1,VCF2[,…],VCFn \
 --out OUTPREFIX
~~~

### 2.3 Statistics and QC utilities

TRTools provides a suite of statistics and QC utilites to allow fast high-level checks of TR runs. `statSTR` allows users to compute locus level statistics on multi-sample TR VCFs, such as the mean allele length, allele frequency distributions, and call rate. It outputs a tab delimited file listing user-specified statistics for each TR. `statSTR` additionally allows outputing plots of allele frequency distributions at specific TRs (Fig. 1a).

**Figure 1:**
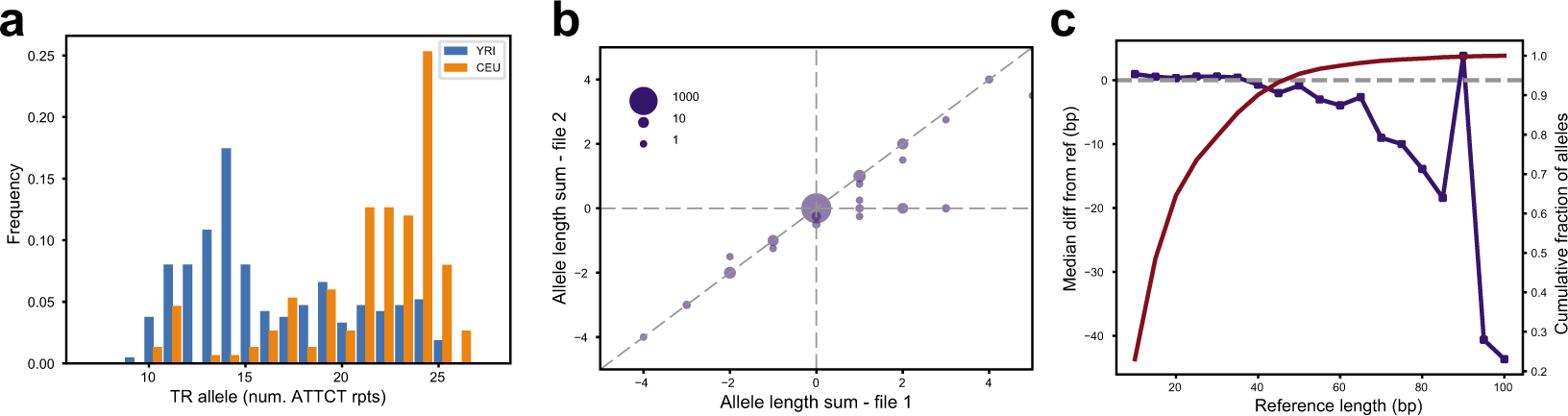
TRTools visualizations. **(a)** Example allele frequency distribution at an example pentanucleotide TR output by statSTR based on GangSTR genotypes for two sample sets (YRI population consisting of Yorubans from Nigeria and CEU population of Northwestern European descent). **(b)** Example TR genotype comparison output by compareSTR. The plot compares genotypes (in terms of number of repeats difference from hg19) from HipSTR (x-axis) to those from ExpansionHunter (y-axis) on 5,000 tetranucleotide TRs. Bubble sizes give the number of calls included in each point. **(c)** Example reference bias plot output by qcSTR using popSTR2 genotypes. The plot shows the average deviation of TR alleles called vs. the reference length of the TR (in bp). The red line shows the cumulative percentage of allele calls below each reference length threshold.

~~~
statSTR --vcf VCF --out OUTPREFIX [statistics]
~~~

`compareSTR` allows users to compare calls from two VCF files. These can be generated by the same or different tools. This allows users to compare calls across platforms and for different runtime options. Fig. 1b shows an example plot created by `compareSTR` comparing two call sets.

~~~
compareSTR --vcf1 VCF1 --vcf2 VCF2 \
   [--vcftype1 VCFTYPE] [--vcftype2 VCFTYPE] \
   --out OUTPREFIX [options] \
~~~

`qcSTR` automatically generates plots for performing quality control of TR genotype datasets. For example, Fig. 1c shows a plot demonstrating an expected deletion bias at long alleles based on popSTR2 genotypes.

~~~
qcSTR --vcf VCF --out OUTPREFIX [options]
~~~

### 2.4 Python library for data analysis

To enable researchers to leverage TRTools features in their own custom tools, we have packaged it as a Python library. The underlying functionality for operations such as harmonizing VCF records across TR genotypers or performing string manipulations on TR sequences can be accessed by importing the library into a Python script.

~~~
import vcf, trtools.utils.tr_harmonizer as trh
reader = vcf.Reader(open(“my.vcf”))
vcftype = trh.InferVCFType(reader)
rec = reader.next()
trrecord = trh.HarmonizeRecord(vcftype, rec)
trrecord.GetAlleleFrequencies(uselength=True)
# {10: 0.2, 15: 0.8} dict of num. rpts.->freq
~~~

**Table 1:**
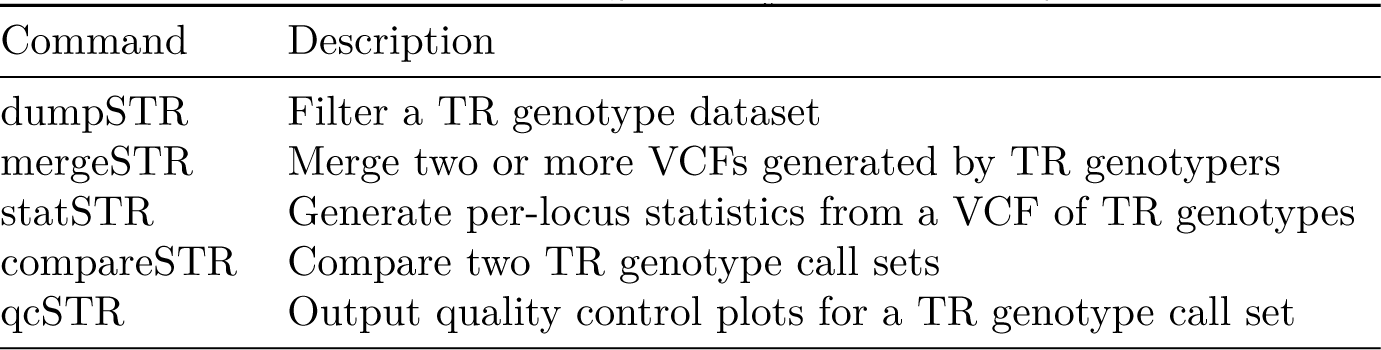
Summary of current TRTools utilities Command Description.

## 3 Discussion

Quality control and filtering are crucial steps for nearly any genome- or population-scale analysis. TRTools meets a pressing need for standardized tools for performing these tasks on TR datasets, which are not handled well by mainstream tools. This toolkit currently supports five major TR genotypers. It can easily be extended to additional tools compatible with VCF standards and to incorporate additional utilities as the community continues to develop standards for TR analysis.

## Acknowledgements

We thank Vineet Bafna, Mehrdad Bakhtiari, Jonghun Park, and Snædis Kristmundsdottir for helpful comments and sharing VCF files. Whole genome sequencing data for individual NA12881 was obtained from the dbGaP (phs001224.v1.p1).

