## Supplementary Material for "TRTools: a toolkit for genome-wide analysis of tandem repeats"

### Supplementary Material - TRTools

#### Datasets

BAM files containing reads from high-coverage whole genome sequencing datasets for the 1000 Genomes Project (Consortium *et al.*, 2015) were accessed through the European Nucleotide Archive accession number `PRJEB31736`. They were processed using GangSTR (Mousavi *et al.*, 2019) v2.4.2.12 with non-default parameter `--grid-threshold 250` using the TR reference file `hg38_ver17.bed.gz` available on the GangSTR website (https://github.com/gymreklab/gangstr). Allele frequencies for a pentanucleotide repeat in the promoter of *RUNX1* (hg38 chr21:35348646-35348646) for samples from the YRI (Yoruba in Ibadan, Nigeria) and CEU (Utah Residents with Northern and Western European Ancestry) populations are shown in Fig. 1a in the main text.

Whole genome sequencing (BAM file aligned to hg19) for Platinum Genomes sample NA12881 was downloaded from dbGaP (accession phs001224.v1.p1). A single chromosome (chromosome 10) was extracted using samtools (Li *et al.*, 2009). TRs were genotyped using ExpansionHunter (Dolzhenko *et al.*, 2017) v3.2.0 and HipSTR (Willems *et al.*, 2017) v0.6.2. For both, we used the GangSTR version 16 reference subsetted to the first 5,000 tetranucleotides as the input set of TRs. ExpansionHunter was run with parameter `-a path-aligner`. HipSTR was otherwise run with default parameters. We used dumpSTR to filter each callset and compareSTR to generate the bubble plot in Fig. 1b using the options shown below

A VCF file generated by popSTR2 (Kristmundsdottir *et al.*, 2019) on Platinum Genomes samples NA12891, NA12892, and NA12878 (dbGaP phs001224.v1.p1) was obtained from the PopSTR authors. This file was used to generate the reference bias plot shown in Fig. 1c in the main text.

#### Commands for generating figures

The following code snippets show the commands used to generate the figures in the main text.

Listing 1: Code to generate Fig. 1a

```
#!/bin/bash
# YRIVCF and CEUVCF were generated by GangSTR v2.4.2.12
REGION=chr21:35348646-35348646
tabix --print-header $YRIVCF $REGION | bgzip -c > yri_runx1.vcf.gz
tabix --print-header $CEUVCF $REGION | bgzip -c > ceu_runx1.vcf.gz
tabix -p vcf yri_runx1.vcf.gz
tabix -p vcf ceu_runx1.vcf.gz

# Merge
mergeSTR --vcfs yri_runx1.vcf.gz,ceu_runx1.vcf.gz --out yri_ceu_runx1
bgzip -f yri_ceu_runx1.vcf
tabix -p vcf -f yri_ceu_runx1.vcf.gz

# Get sample lists
bcftools query -l yri_runx1.vcf.gz > yri_samples.txt
bcftools query -l ceu_runx1.vcf.gz > ceu_samples.txt

# StatSTR
# Compute stats separately on YRI and CEU samples
statSTR \
    --vcf yri_ceu_runx1.vcf.gz \
    --samples yri_samples.txt,ceu_samples.txt --sample-prefixes YRI,CEU \
    --region $REGION \
    --out yri_ceu_runx1 \
    --afreq --use-length --plot-afreq
# Output file yri_ceu_runx1-chr21-35348646.pdf shown in Fig. 1a
```

Listing 2: Code to generate Fig. 1b

```
#!/bin/bash

SAMPLE=NA12881
# $SAMPLE-hipstr.vcf.gz and $SAMPLE-eh-path.vcf.gz generated
# by calling HipSTR and ExpansionHunter on the same TR reference

# Filter
dumpSTR \
    --vcf $SAMPLE-hipstr.vcf.gz \
    --hipstr-min-call-Q 0.9 \
    --hipstr-min-call-DP 10 \
    --hipstr-max-call-DP 1000 \
    --hipstr-max-call-flank-indel 0.15 \
    --hipstr-max-call-stutter 0.15 \
    --hipstr-min-supp-reads 2 \
    --out $SAMPLE-hipstr.filtered
cat $SAMPLE-hipstr.filtered.vcf | vcf-sort | \
        bgzip -c > $SAMPLE-hipstr.filtered.vcf.gz
tabix -p vcf $SAMPLE-hipstr.filtered.vcf.gz

dumpSTR \
    --vcf $SAMPLE-EH-path.vcf \
    --vcftype eh \
    --eh-min-call-LC 50 \
    --out $SAMPLE-eh-path.filtered
# Edit sample name to be same
cat $SAMPLE-eh-path.filtered.vcf | sed 's/NA12881.chr10/NA12881/' | \
        vcf-sort | bgzip -c > $SAMPLE-eh-path.filtered.vcf.gz
tabix -p vcf $SAMPLE-eh-path.filtered.vcf.gz

# Add contigs to EH
zcat $SAMPLE-hipstr.filtered.vcf.gz | grep congi > hg19_contigs.txt
bcftools annotate -h hg19_contigs.txt $SAMPLE-eh-path.filtered.vcf.gz | \
        bgzip -c > $SAMPLE-eh-path-contigs.filtered.vcf.gz
tabix -p vcf -f $SAMPLE-eh-path-contigs.filtered.vcf.gz

# Make bubbles plot and compare
compareSTR \
    --vcf1 $SAMPLE-eh-path-contigs.filtered.vcf.gz \
    --vcf2 $SAMPLE-hipstr.filtered.vcf.gz \
    --vcftype1 eh \
    --vcftype2 hipstr \
    --out eh-path-hipstr \
    --bubble-min -5 --bubble-max 5
# Output file eh-path-hipstr-bubble-periodALL.pdf shown in Fig. 1b
```

Listing 3: Code to generate Fig. 1c

```
#!/bin/bash
qcSTR --vcf $popSTR_vcf --out popstr_qc
# Output file popstr_qc-diffref-bias.pdf shown in Fig. 1c
```
